# Cornichon Homolog-3 (CNIH3) Modulates Spatial Memory in Female Mice

**DOI:** 10.1101/724104

**Authors:** Hannah E. Frye, Sidney B. Williams, Christopher R. Trousdale, Elliot C. Nelson, Joseph D. Dougherty, Jose A. Morón

## Abstract

Cornichon homolog-3 (CNIH3) is an AMPA receptor (AMPAR) auxiliary protein that traffics AMPARs to the postsynaptic membrane and potentiates AMPAR signaling. AMPARs are key components of hippocampal synaptic plasticity and memory formation, however the role of CNIH3 in memory has yet to be elucidated. To study the role of CNIH3 on mouse behavior, we bred and characterized a line of *Cnih3*^-/-^ mice from C57BL/6 *Cnih3^tm1a(KOMP)Wtsi^* mice obtained from the Knockout Mouse Project (KOMP). In agreement with previous studies of CNIH3 in the brain, we observed concentrated expression of *Cnih3* in the dorsal hippocampus, a region associated with spatial learning and memory. Therefore, we tested *Cnih3*^+/+^, *Cnih3*^+/-^, and *Cnih3*^-/-^ mice in the Barnes maze paradigm to measure spatial memory. We observed no change in spatial memory in male *Cnih3*^+/-^ and *Cnih3*^-/-^ mice compared to male *Cnih3*^+/+^ controls, however, *Cnih3*^-/-^ female mice made significantly more primary errors, had a higher primary latency, and took less efficient routes to the target in the maze compared to *Cnih3*^+/+^ female mice. Next, to investigate an enhancement of spatial memory by *Cnih3* overexpression, specifically in the dorsal hippocampus, we developed an AAV5 viral construct to express wild-type *Cnih3* in excitatory neurons. Female mice overexpressing *Cnih3* made significantly fewer errors, had a lower primary latency to the target, and took more efficient routes to the maze target compared to YFP expressing control females. No change in spatial memory was observed in male *Cnih3* overexpression mice. This study, the first to identify sex-specific effects of the AMPAR auxiliary protein CNIH3 on spatial memory, provides the groundwork for future studies investigating the role of CNIH3 on sexually dimorphic AMPAR-dependent behavior and hippocampal synaptic plasticity.

## INTRODUCTION

α-amino-3-hydroxy-5-methyl-4-isoxazolepropionic acid receptors (AMPARs) are glutamatergic neurotransmitter receptors found abundantly throughout the brain which mediate a wide range of neural processes such as synaptic plasticity, learning and memory behaviors (Cheng et al., 2012). In particular, AMPAR activity in the hippocampus plays a key role in spatial memory (Lee et al., 2003; Matsuo et al., 2008), and AMPAR auxiliary proteins are critical actors underlying AMPAR-dependent learning and memory mechanisms (Volk et al., 2010; Gandhi et al., 2014; Li et al., 2017). Cornichon homolog (CNIH) proteins are an important class of auxiliary proteins for AMPARs in the brain. Cornichon homologs 2 and 3 (CNIH2 and CNIH3) function as AMPAR chaperones from the endoplasmic reticulum (ER) and golgi to the post-synaptic density (PSD) (Shi et al., 2010; Harmel et al., 2012; Brockie et al., 2013). At the postsynaptic membrane, CNIH proteins also act to potentiate AMPAR glutamate sensitivity (Coombs et al., 2012; Haering et al., 2014), improve ion channel permeability (Coombs et al., 2012; Brown et al., 2018), reduce decay of AMPAR excitatory post-synaptic currents (EPSCs) (Boudkkazi et al., 2014), and slow receptor deactivation by delaying internalization (Coombs et al., 2012; Mauric et al., 2013; Shanks et al., 2014; Challenor et al., 2015).

However, little is known about the effect of CNIH proteins on AMPAR-associated mammalian behavior or pathology. A correlative study of post-mortem human brain tissue found abnormal expression of CNIH1, CNIH2, and CNIH3 in the brains of patients diagnosed with schizophrenia compared to the brains of non-schizophrenic patients (Drummond et al., 2012). A 2012 case study reported a deletion of *CNIH2* in a patient with dysmorphic features and intellectual disability (Floor et al., 2012). Recently, we and our colleagues performed a genome-wide association study (GWAS) that found several single-nucleotide polymorphisms (SNPs) in the noncoding region of human *CNIH3* significantly associated with decreased risk for opioid dependence (Nelson et al., 2016). We have also reported the significance of AMPARs in opioid-associated memory in mice (Morón et al., 2007; Billa et al., 2009; Billa et al., 2010; Xia et al., 2011), therefore we hypothesized a potential link between the AMPAR auxiliary protein CNIH3 and memory.

AMPAR-mediated currents, receptor subunit configuration, post-translational modifications, synaptic membrane localization, and protein-protein interactions shape a wide range of learning and memory behaviors (Lee et al., 2003; Matsuo et al., 2008; Sanderson et al., 2008; Penn et al., 2017). For example, the transmembrane AMPAR regulatory protein (TARP) stargazin is upregulated in the cerebellum following eye-blink conditioning in rats (Kim and Thompson, 2011). PSD-95, an excitatory synaptic scaffolding protein which aids in maintaining AMPARs at the synaptic membrane, plays a role in spatial memory, fear conditioning, and extinguished memory retrieval (Nagura et al., 2012; Gandhi et al., 2014; Li et al., 2017). Pick1 is involved in AMPAR removal from the synaptic membrane and is necessary for inhibitory avoidance memory (Volk et al., 2010). Post-translational modifications of these proteins, such as the phosphorylation of the TARP γ-8 by CAMKII is necessary for both context and cue-associated fear conditioning in mice (Park et al., 2016). Therefore, we hypothesize that CNIH3 similarly mediates AMPAR activity underlying memory processes in the hippocampus.

In this study, we developed and characterized a line of *Cnih3* knockout (KO) mice to study the role of CNIH3 on spatial memory. We found high CNIH3 expression in the dorsal hippocampus, a brain region where AMPAR activity plays a key role in the modulation of spatial memory (Tzakis et al., 2016; Torquatto et al., 2019). Therefore, we hypothesize that spatial memory is impaired in *Cnih3*^-/-^ mice compared to *Cnih3*^+/+^ age-matched controls. Interestingly, we observed a distinct effect of *Cnih3* expression on mouse performance in the Barnes maze spatial memory task dependent on the sex of the animal. We found that both global KO of *Cnih3* and hippocampal overexpression of *Cnih3* affect spatial memory only in female animals. This study builds on previous literature which found CNIH3 to play a key role in the maintenance of AMPAR signaling and function and adds new knowledge about how sex may play a role in AMPAR-associated spatial memory.

## MATERIALS AND METHODS

### Animals

All experimental protocols utilizing animals were approved by the Institutional Animal Care and Use Committee at Washington University in St. Louis. Male and female mice between 8-12 weeks of age were used for all experiments. Breeding for the *Cnih3^tm1a(KOMP)Wtsi^* and the *Cnih3*^-/-^ mouse colonies are described in the Experimental Results and in Figure 1B.

**Figure 1:**
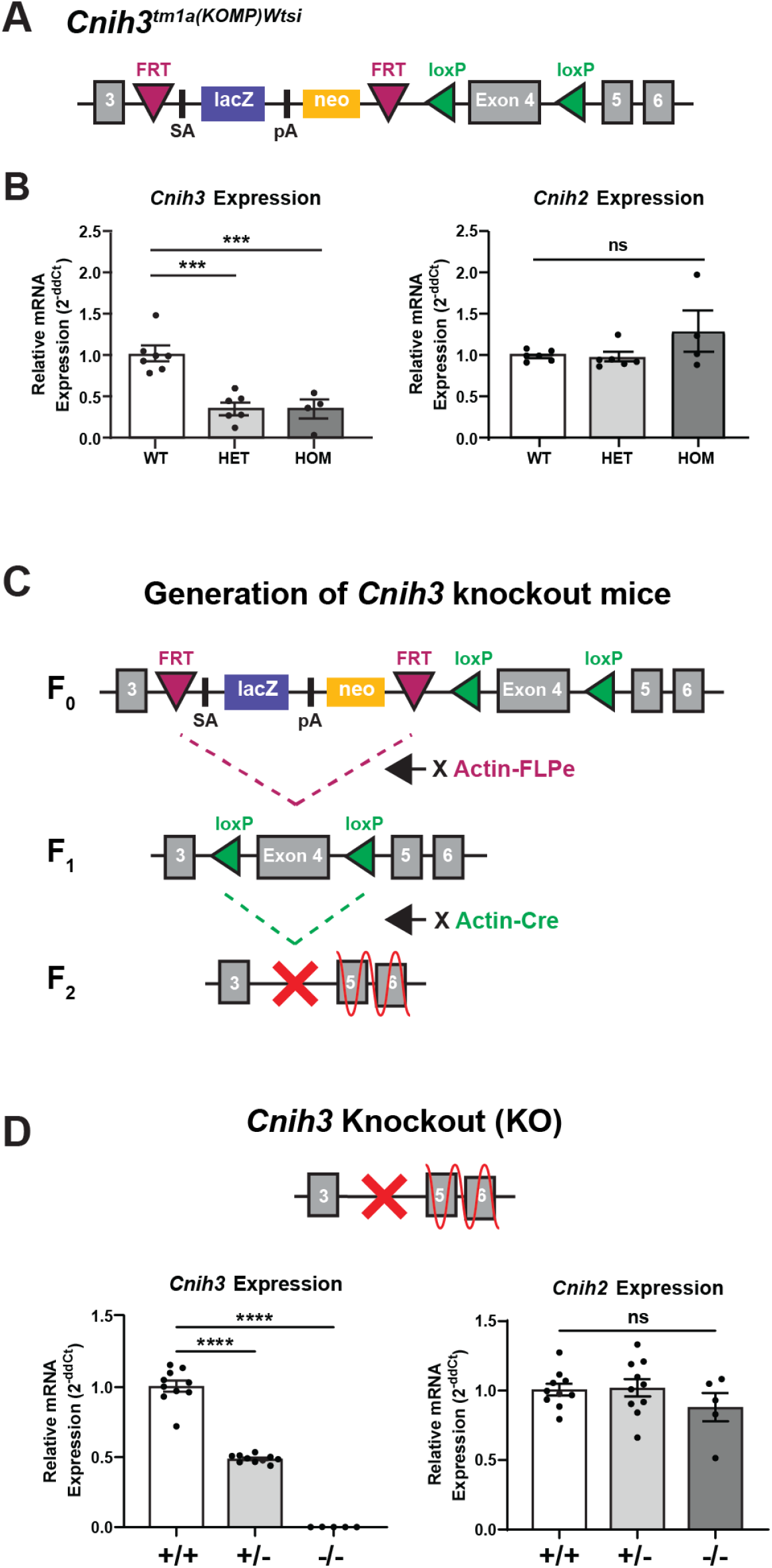
Generation and characterization of *Cnih3* knockout (KO) mice. (A) The KOMP *Cnih3^tm1a(KOMP)Wtsi^* gene contains a polyadenylation (pA) site after exon 3 to halt mRNA transcription. The pA site as well as *lacZ* and *neo* cassettes and a splice acceptor (SA) site are contained within two *FRT* sites. Exon 4 is surrounded by two *loxP* sites *Cnih3^tm1a(KOMP)Wtsi^* mice were bred to create a colony of WT, HET, and HOM *Cnih3^tm1a(KOMP)Wtsi^* mice. (B) RT-qPCR results for *Cnih3^tm1a(KOMP)Wtsi^* WT (n = 7), HET (n = 6), and HOM (n = 4) mice for expression of *Cnih3* and *Cnih2* mRNA. Relative mRNA expression is presented as relative fold change calculated by 2^-(ddCt)^. (C) Breeding scheme for *Cnih3* KO mice from *Cnih3^tm1a(KOMP)Wtsi^* mice. (D) RT-qPCR results for *Cnih3* and *Cnih2* mRNA expression in *Cnih3*^+/+^ (n = 10), *Cnih3*^+/-^ (n = 10), and *Cnih3*^-/-^ (n = 5) mice (one-way ANOVA for genotype and *post-hoc* Sidak’s multiple comparison test, * denotes significance compared to WT (ns *p* > 0.05, *** *p* < 0.001).

### Real-time quantitative PCR

Mice were euthanized and the dorsal hippocampi from individual mice were rapidly dissected and immediately frozen on dry ice for real-time quantitative PCR (RT-qPCR) analysis. Tissue was homogenized for RNA extraction using the RNeasy Mini Kit (QIAGEN). RNA quantity and quality were measured with a Nanodrop Spectrophotometer (Thermo Fisher Scientific). 500 ng of RNA was used to synthesize cDNA via reverse transcription using iScript Reverse Transcriptase Supermix (Bio-Rad). Each reaction tube contained a volume of 10μL containing 500 nM of forward and reverse primers, 5 μL PowerUp SYBR Green Master Mix (QIAGEN), and 4 μL of purified cDNA diluted 1:2. The reaction was carried out in an Applied Biosystems Quant Studio 6 PCR system (Thermo Fisher Scientific). The Ct scores from each sample were normalized to the expression of β-actin and to wild-type (WT) or viral controls. The expression fold change in gene expression was calculated using Double Delta Ct Analysis (ddCt).

### Primer sequences

Primers used for RT-qPCR were obtained from Integrated DNA Technologies. Primer sequences are as follows: *Cnih3* exon 4: 5’-TGGTGCTGCCCGGAGT-3’ (forward), 5’-CCAGAAGTGATAGAAAAGCAGAG-3’ (reverse). Viral *Cnih3* gene variant 2: 5’-GCGCTGCGCTCATCTTTTTC-3’ (forward), 5’-CTTGAAATCCGTTCTTAGCTCGT-3’ (reverse). *Cnih2*: 5’-ATATTCCATCCACGGCCTCTTCTGTCTGA-3’ (forward), 5’-AGAAGAAGGACAGCAGGTAGAAGGCGAGTTTG-3’ (reverse).

### β-galactosidase staining

To visualize the anatomical distribution of the *Cnih3^tm1a(KOMP)msι^* gene in the brain, we stained brains of homozygous male and female mice using a β-galactosidase staining assay (West et al., 2015; Trifonov et al., 2016) to identify expression of the *lacZ* cassette contained within the *Cnih3^tm1a(KOMP)Wtsi^* gene. Mice were transcardially perfused with ice-cold 4% paraformaldehyde (PFA) in phosphate-buffered saline (PBS) while deeply anesthetized with isofluorane. The whole brain was extracted and post-fixed overnight in 4% PFA before the brains were transferred for equilibration in 30% sucrose in PBS. Equilibrated brains were flash-frozen and the entire brain was sectioned into 40 μm slices using a cryostat. Floating slices were washed 3×15 minutes with rinse buffer (0.1M PB, 2 mM MgCl2, 0.1% Triton X-100, 0.01% Deoxycholic Acid, and 1.25 mM EGTA, pH 7.4) and then placed in X-gal staining solution (1 mg/mL X-gal and 0.4 mg/mL Nitrotetrazolium blue in rinse buffer) for 18 hours overnight at 37 °C. Following overnight incubation, brain slices were rinsed 3×15 minutes in rinse buffer, mounted onto slides, dehydrated in successive washes of ethanol (50, 75, 95, 100, and 100%), cleared in three changes of xylene, and cover-slipped using Cytoseal XYL mounting media (Richard-Allen). Slides were imaged on a Zeiss Axio Scan Z1 Brightfield slide scanner microscope (Zeiss).

### Barnes maze spatial learning task

To assess changes in spatial learning and memory, a Barnes maze spatial learning task was conducted as previously described (Fakira et al., 2016). Mice were handled by the experimenter for three days prior to the experiment to minimize handling stress. The Barnes maze apparatus was a 91 cm circular platform surround by twenty equidistant 5 cm holes. One hole, referred to as the target hole, led to a dark escape box in which the mouse could hide. All other holes led to false bottoms. Spatial cues in the form of large shapes (triangle, square, hexagon, and cross) surrounded the apparatus to allow for spatial navigation (Barnes, 1979). Any-Maze tracking software (Stoelting) was used for video recording and animal tracking. During habituation to the apparatus, the mouse was gently placed in the center of the maze and allowed to explore the maze for 180s in two consecutive trials. Upon entering the target hole, the entrance was covered, and the animal remained in the target box for 1 minute. If the animal did not enter the target hole in the 180s period, the animal was gently guided to the hole. During training days 1-4, three 180s trials were conducted for each animal in 15min intervals. The table was thoroughly cleaned between each trial with 70% ethanol and the top table was rotated to minimize olfactory cues leading to the target hole. The latency to the first entry in the target hole (primary latency), the number of errors made prior to initial target hole entry (primary errors), and the path efficiency (actual distance traveled to target hole/shortest distance to target hole from starting location) were recorded. On Day 5, a 90s probe trial was conducted where the escape box was replaced with a false bottom to prevent entry. The primary latency, primary errors, and path efficiency were recorded.

### Generation of a Cnih3 expressing virus

A viral vector was constructed using pENTR3C and the *mus musculus* mRNA sequence of CNIH3, variant 2 (Genbank Accession: BC115640). Transcriptional variant was chosen based on expression level in the brain and hippocampus, as notated on the UCSC Genome browser (Kent et al., 2002). CNIH3 cDNA was PCR’ed from clone 40103001 from the Mammalian Gene Collection (MGC) and cloned into pENTR3C adding an N-terminal Myc-tag and featured a t2A sequence followed by a GFP sequence to mark expressing cells. Once assembled in pENTR3c, the transcript was transferred into a pAAV-EF1a using Gateway cloning. Upon confirmation of cloning for this vector, the EF1A promoter was replaced by a CamKIIa promoter via classical restriction enzyme-based cloning, resulting in the pAAV-CAMKII-myc-CNIH3-t2a-GFP construct. The plasmid, once fully sequence verified, was packaged with an AAV5 serotype at the Hope Center Viral Core at Washington University in St Louis.

### Intracranial injection surgeries

WT mice were anesthetized with 1-3% isofluorane and head-fixed in a stereotaxic apparatus (Stoelting). 0.5 μL of AAV5-CAMKII-myc-CNIH3-t2a-GFP virus (1.7×10^13^ particles per mL) or 0.5 μL of AAV5-CAMKII-eYFP (8.6×10^11^ particles per mL, Virus Vector Core, The University of North Carolina at Chapel Hill) control virus was slowly infused bilaterally into the dorsal hippocampus (A/P: −1.8, Lat: ±1.4, D/V: −1.8). After three weeks, hippocampi were either extracted for RT-qPCR or mice underwent training in the Barnes maze. Following Barnes maze testing, mice were transcardially perfused with 4% PFA and brains were sectioned for immunohistochemical verification of viral placement.

### Immunohistochemistry

40 μm free-floating brain slices from viral *Cnih3* overexpression mice were stained with rabbit anti-myc primary antibody (Cell Signaling Technology, #2272) and goat anti-rabbit 594 Alexa Fluor secondary antibody (Life Technologies, #A11037). Slices were mounted and imaged via confocal microscope to visualize virally expressed myc-tagged CNIH3 protein (Figure 4A).

### Statistics and data analysis

For statistical analysis, datasets were analyzed using GraphPad Prism 8 software. For RT-qPCR experiments, one-way ANOVAs and unpaired two-tailed t tests were used to compare mRNA expression between genotypes and viral overexpression compared to WT and YFP expressing controls. Normality of each dataset was determined using a Shapiro-Wilk test. After determining that male and female mice did not differ in gene expression, data were pooled across sexes. For behavioral analysis, datasets were analyzed by two-way ANOVA for genotype and sex, with *post-hoc* Sidak’s multiple comparison tests used to compare genotypes and viral overexpressing animals to WT and viral controls. One female *Cnih3*^+/-^ mouse froze during the probe trial and did not move during the test; therefore, it was eliminated from the final analysis. Outlier values were identified using a Grubbs’ test and animals which were outliers in primary errors, primary latency, or path efficiency datasets were eliminated from the final analysis (one male and female *Cnih3*^+/+^ and *Cnih3*^+/-^, one male *Cnih3*^-/-^ mouse, one female overexpressing mouse). In addition, viral expression and injection placement were confirmed for every animal which underwent intracranial viral injection prior to behavioral testing, and one female injected with YFP control virus was excluded from final analysis due to failure of viral expression. The normality of each dataset was determined using a Shapiro-Wilk test. *Cnih3*^+/+^ and *Cnih3*^+/-^ datasets did not pass normality for primary errors and the *Cnih3* overexpressing female dataset did not pass normality for path efficiency in the Barnes maze, therefore these datasets were also analyzed using a nonparametric Kruskal-Wallis or Mann-Whitney test within each sex. Changes in gene expression and spatial memory parameters were considered statistically significant with *p* < 0.05.

## EXPERIMENTAL RESULTS

### Generation and validation of a Cnih3 knockout (KO) mouse line

To examine the involvement of CNIH3 in learning and memory behaviors, we developed and characterized a line of *Cnih3*^+/+^, *Cnih3*^+/-^, and *Cnih3*^-/-^ C57BL/6 mice bred from *Cnih3^tm1a(KOMP)Wtsi^* mice. *Cnih3^tm1a(KOMP)Wtsi^* heterozygous male C57BL/6N mice were obtained from the Knockout Mouse Project (KOMP) (IMPC, 2016) and backcrossed with WT female C57BL/6J mice to produce *Cnih3^tm1a(KOMP)Wtsi^* heterozygous offspring on a C57BL/6J background. Heterozygous *Cnih3^tm1a(KOMP)Wtsi^* male and female mice were bred to create WT, heterozygous (HET), and homozygous (HOM) *Cnih3^tm1a(KOMP)Wtsi^* offspring. The *Cnih3^tm1a(KOMP)Wtsi^* gene contained a polyadenylation (pA) site following exon 3 to attenuate transcription of subsequent exons and was thus designed to be a “knockout-first” allele (Figure 1A). However, RT-qPCR analysis of mRNA synthesized cDNA determined that *Cnih3^tm1a(KOMP)Wtsi^* heterozygote and homozygote animals expressed only a 60% reduction in exon 4 of *Cnih3* compared to WT mice (One-way ANOVA, F_(2,14)_ = 20.40, *p* < 0.0001, *post-hoc* Sidak’s multiple comparisons test to WT, *p* = 0.0001 and 0.0003, respectively), resulting in a *Cnih3* knockdown (KD) instead of a full KO animal (Figure 1A). As CNIH2 is a functionally similar homolog of CNIH3, we also conducted RT-qPCR to assess *Cnih2* expression to determine if *Cnih3* KD results in compensatory expression of *Cnih2*. No change in *Cnih2* expression was observed in *Cnih3^tm1a(KOMP)Wtsi^* hetero- or homozygote animals compared to WT controls (One-way ANOVA, F_(2,13)_ = 2.041, *p* = 0.1695) (Figure 1A). Male and female mice did not differ in gene expression levels.

To create a line of total *Cnih3* KO animals, we conducted additional breeding to eliminate exon 4 from the *Cnih3^tm1a(KOMP)Wtsi^* gene (Figure 1B). *Cnih3^tm1a(KOMP)Wtsi^* homozygote males were bred with Actin-FLPe females to excise the Splice Acceptor site (SA), pA site, and the *lacZ* and *neomycin* (*neo*) cassettes contained within the *FRT* sites (generation F1). Male F1 mice were bred with Actin-Cre females to excise exon 4 contained with the *loxP* sites (generation F2). Removal of exon 4 results in a frameshift mutation across exons 5 and 6 which results in nonsense mediated decay and complete loss of function. RT-qPCR analysis found a significant decrease in *Cnih3* expression between genotypes (one-way ANOVA, F_(2,22)_ = 264.1, *p* < 0.0001). *Cnih3*^+/-^ mice expressed 50% of *Cnih3* exon 4 compared to WT animals and *Cnih3*^-/-^ mice have a total elimination of *Cnih3* exon 4 expression (*post-hoc* Sidak’s multiple comparisons test, *p* < 0.0001 each genotype) (Figure 1C). *Cnih2* expression remained unchanged in *Cnih3*^+/-^ and *Cnih3*^-/-^ mice compared to WT mice (one-way ANOVA, F_(2,22)_ = 1.097, *p* = 0.3513). No difference was observed in gene expression between male and female mice.

### Anatomical expression of Cnih3 in the brain

To identify where *Cnih3* was expressed in the brain, we visualized *Cnih3* expression using a β-galactosidase staining assay to probe the *lacZ* cassette contained within the *Cnih3^tm1a(KOMP)Wtsi^* gene (Figure 2). *Cnih3* expression was highest in the prefrontal cortex (PFC), hypothalamus, cortex, amygdala, and hippocampus. Within the hippocampus, a region where AMPAR activity regulates spatial and contextual memory (Lee et al., 2003; Matsuo et al., 2008; Sanderson et al., 2008; Xia et al., 2011; Sebastian et al., 2013; Penn et al., 2017), *Cnih3* expression was especially pronounced. In particular, the highest *Cnih3* expression was observed within the dentate gyrus (DG) and the CA1 regions of the dorsal hippocampus (Figure 2). No difference in *Cnih3* expression was observed between male and female mice. While confirmation of null CNIH3 protein expression would be ideal to validate *Cnih3*^-/-^ animals and to probe for anatomical expression of CNIH3, we were unable to obtain or validate a suitable antibody for CNIH3. Nonetheless, given that our Lacz study concurred with previous studies showing high expression of cornichon proteins in the hippocampus (Kato et al., 2010; Herring et al., 2013), we next examined spatial memory in these mice.

**Figure 2:**
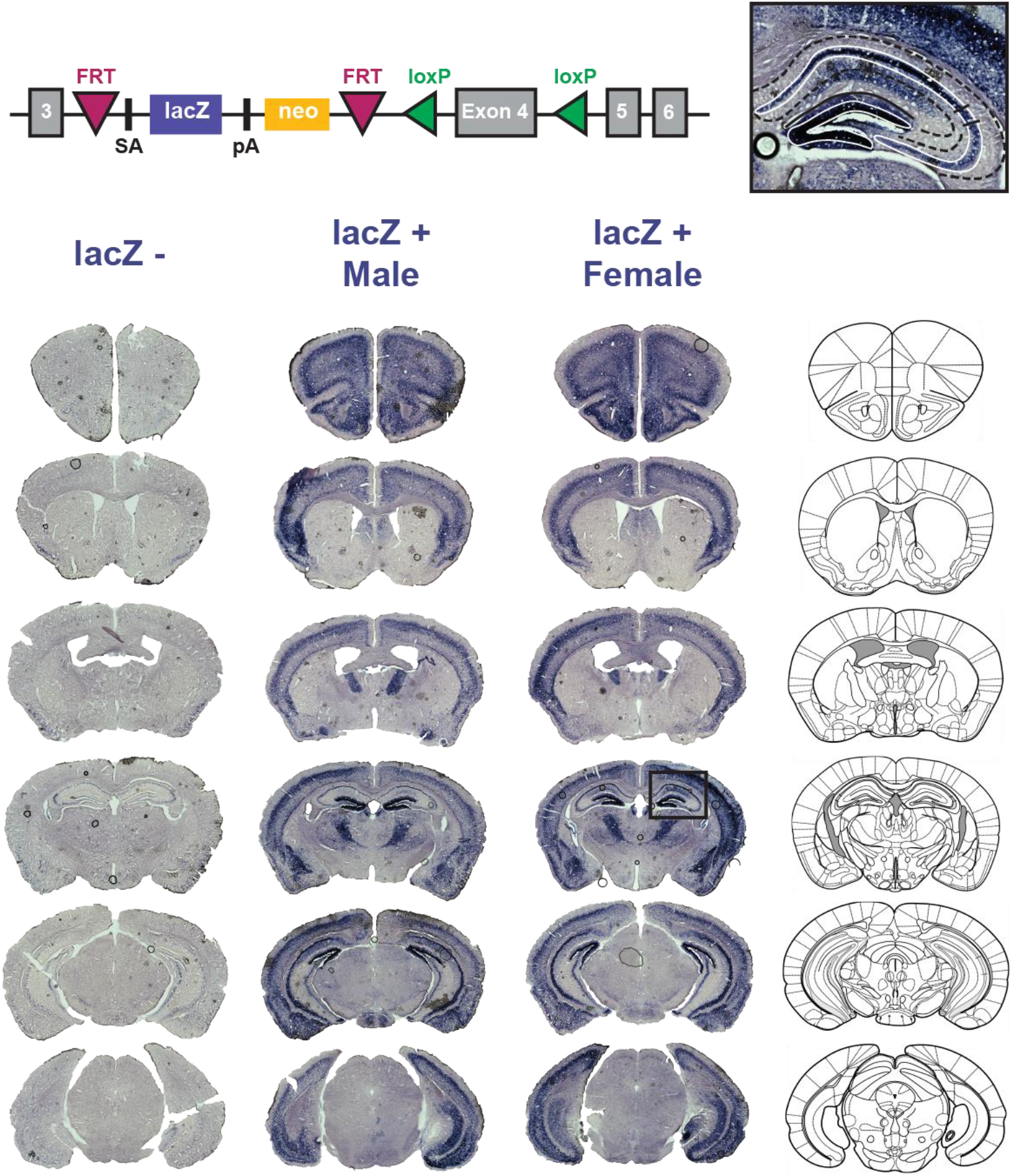
*Cnih3* is strongly expressed in the dorsal hippocampus of *Cnih3^tm1a(KOMP)Wtsi^* mice. β-galactosidase staining for the *lacZ* cassette contained within the *Cnih3^tm1a(KOMP)Wtsi^* gene qualitatively identifies *Cnih3* expression throughout the brain. Representative slices from *lacZ*− and from *lacZ*+ male and female are shown. A closeup image of a *lacZ*+ dorsal hippocampus shows strong *lacZ* expression in the CA1 and dentate gyrus regions of the dorsal hippocampus.

### Spatial memory is impaired in Cnih3 KO female mice

AMPAR activity in the hippocampus plays a critical role in spatial memory (Lee et al., 2003; Matsuo et al., 2008), and AMPAR auxiliary proteins facilitate AMPAR-dependent mechanisms that underlie these behaviors (Volk et al., 2010; Gandhi et al., 2014; Li et al., 2017). To determine if CNIH3 plays a role spatial memory, we utilized the Barnes maze behavioral paradigm (Barnes,1979; Sunyer et al., 2007) to assess spatial memory in *Cnih3*^+/-^ and *Cnih3*^-/-^ mice (Figure 3A). In the Barnes maze, mice were trained to use spatial cues around the room to locate a target hole on a large circular table leading to a dark box in which the mouse could hide. During the Day 5 probe, we observed significant main effects of sex and genotype on animal performance in primary errors (two-way ANOVA, sex [F_(1, 54)_ = 12.58, *p* = 0.0008], genotype [F_(2, 54)_ = 8.539, *p* = 0.0006], and interaction [F_(2, 54)_ = 12.60, *p* < 0.0001]) (Figure 3B) and primary latency (sex [F_(1, 54)_ = 16.69, *p* = 0.0001], genotype [F_(2, 54)_ = 7.549, *p* = 0.0013], and interaction [F_(2, 54)_ = 15.03, *p* < 0.0001]) (Figure 3C), and a significant effect of sex on path efficiency (sex [F_(1, 54)_ = 13.85, *p* = 0.0005], genotype [F_(2, 54)_ = 1.690, *p* = 0.1941], and interaction [F_(2, 54)_ = 7.870, *p* = 0.0010]) (Figure 3D). The sex-dependent effect of genotype on primary errors in the Barnes maze was also confirmed using a nonparametric Kruskal-Wallis test due to the presence of non-normal datasets (Female mice, *p* = 0.0065; Male mice, *p* = 0.6160). Overall, we conclude that *Cnih3* expression significantly modulates performance in the Barnes maze in a sex-specific manner.

**Figure 3:**
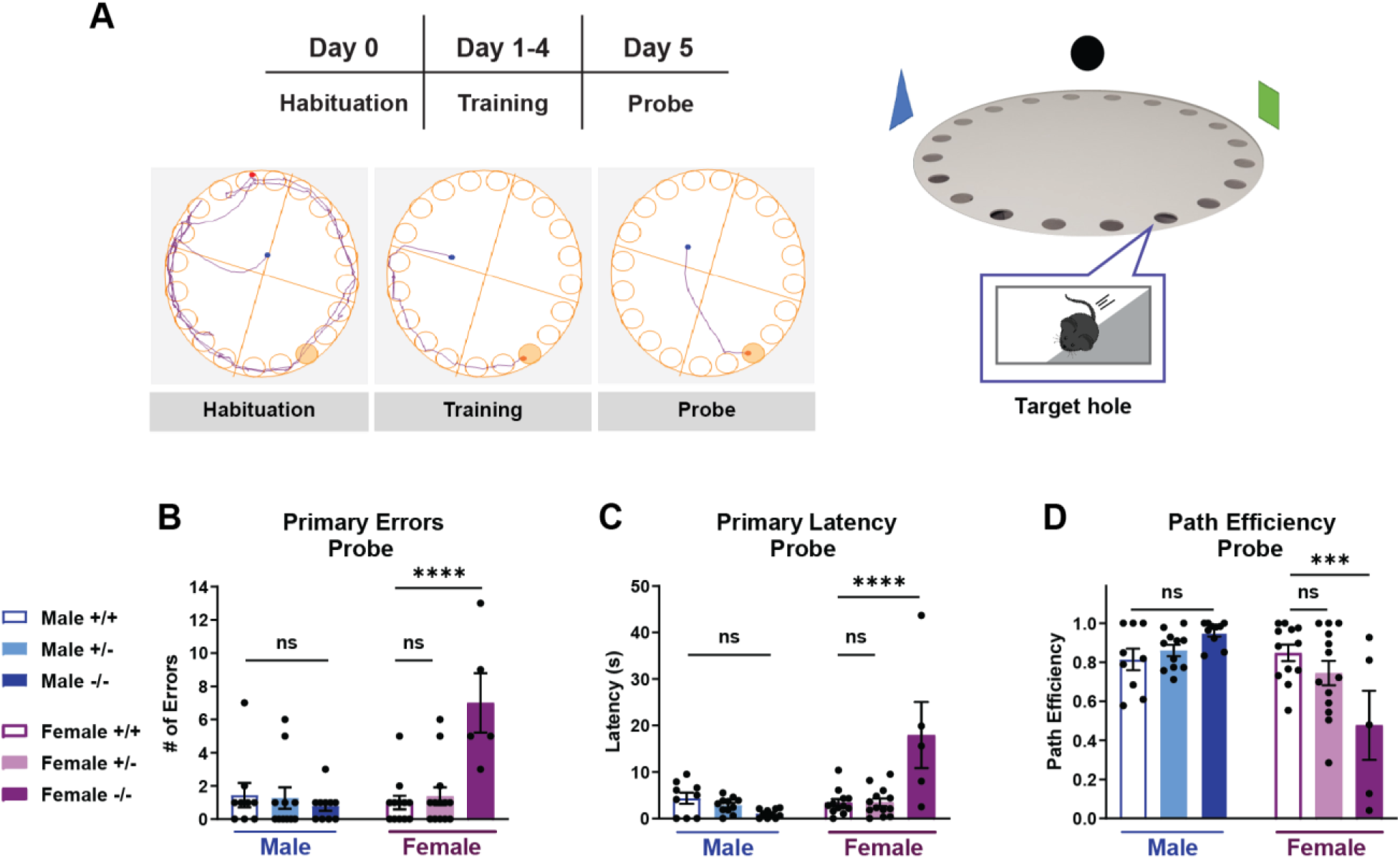
Spatial memory is impaired in *Cnih3*^-/-^ female mice, but not male mice. (A) Schematic of the Barnes maze protocol used to measure spatial memory in mice. (B – D) Barnes maze results of male and female *Cnih3*^+/+^ (male: n = 9; female: n = 12), *Cnih3*^+/-^ (male: n = 10; female: n = 13), and *Cnih3*^-/-^ (male: n =10; female: n = 5) mice during the probe trails is represented by (B) primary errors before location of the target hole, (C) primary latency to entry into the target hole, and (D) path efficiency to the target hole (two-way ANOVA for genotype and sex and *post-hoc* Sidak’s multiple comparisons test (ns *p* > 0.05, *** *p* < 0.001, **** *p* < 0.0001).

To further dissect the interaction between sex and genotype on spatial memory in the Barnes maze, *post-hoc* multiple comparisons were performed to compare *Cnih3*^+/-^ and *Cnih3*^-/-^ to sex-matched WT controls. We found that female *Cnih3*^-/-^ mice made significantly more primary errors (*post-hoc* Sidak’s multiple comparisons test, *p* < 0.0001; nonparametric Dunn’s multiple comparison’s test, *p* = 0.0057) (Figure 3B), were slower to reach the target (*post-hoc* Sidak’s multiple comparisons test, *p* < 0.0001) (Figure 3C), and took less efficient paths to the target (*p* = 0.0005) (Figure 3D) compared to WT female mice. However, no significant changes in primary errors (*post-hoc* Sidak’s multiple comparisons test, *p* = 0.9769; nonparametric Dunn’s multiple comparison’s test, *p* > 0.9999), primary latency (*post-hoc* Sidak’s multiple comparisons test, *p* = 0.9904), nor path efficiency (*p* = 0.2547) were observed in *Cnih3*^+/-^ female mice compared to female WT mice (Figure 3B-D). Conversely, male mice displayed no significant difference in the number of primary errors (*post-hoc* Sidak’s multiple comparisons test, *Cnih3*^+/-^: *p* = 0.9417 and *Cnih3*^-/-^: 0.74444; nonparametric Dunn’s multiple comparison’s test, *Cnih3^+/-^*: *p* = 0.6501 and *Cnih3^-/-^*: *p* > 0.9999) (Figure 3B), primary latency to the target (*post-hoc* Sidak’s multiple comparisons test, *Cnih3^+/-^*: *p* = 0.7503 and *Cnih3^-/-^*: *p* = 0.2784) (Figure 3C), nor path efficiency (*Cnih3^+/-^*: *p* = 0.8664 and *Cnih3^-/-^*: *p* = 0.1967) (Figure 3D). We did not find evidence for sex differences in the performance of WT animals in the Barnes maze; male and female WT mice did not exhibit differences in primary errors (*p =* 0.9931), primary latency (*p =* 0.9875), nor path efficiency (*p =* 0.9514). Our results demonstrate that CNIH3 is necessary for spatial learning and memory in a sex-dependent manner.

### Validation of a new Cnih3 viral overexpression construct

Since *Cnih3* expression is necessary for spatial memory in female mice, therefore we wanted to determine if supraphysiological hippocampal *Cnih3* is sufficient to enhance spatial memory. In order to investigate this, we generated an AAV5-CAMKII-myc-CNIH3-t2a-GFP virus to induce local overexpression of *Cnih3* in hippocampal excitatory neurons. Bilateral injections of AAV5-CAMKII-myc-CNIH3-t2a-GFP or AAV5-CAMKII-eYFP control virus were performed, targeting the dorsal hippocampus in WT animals to induce *Cnih3* overexpression (Figure 4A). To verify overexpression of *Cnih3*, RT-qPCR was performed to measure mRNA expression of *Cnih3* and *Cnih2* (Figure 4B). The dorsal hippocampus of mice injected with the *Cnih3* overexpression virus expressed ~200X more *Cnih3* mRNA compared to YFP expressing controls (two-tailed unpaired *t*-test, *p* < 0.0001). Viral overexpression of *Cnih3* did not result in a compensatory change in *Cnih2* expression in the dorsal hippocampus (two-tailed unpaired *t*-test, *p* = 0.0593). Injection location and spread of virus were verified through immunofluorescent probing for the myc-tag adjacent to CNIH3 (Figure 4C). This new tool allowed us to overexpress *Cnih3* only in the dorsal hippocampus to measure the consequences of CNIH3 overexpression specifically in this region on spatial memory behavior.

**Figure 4:**
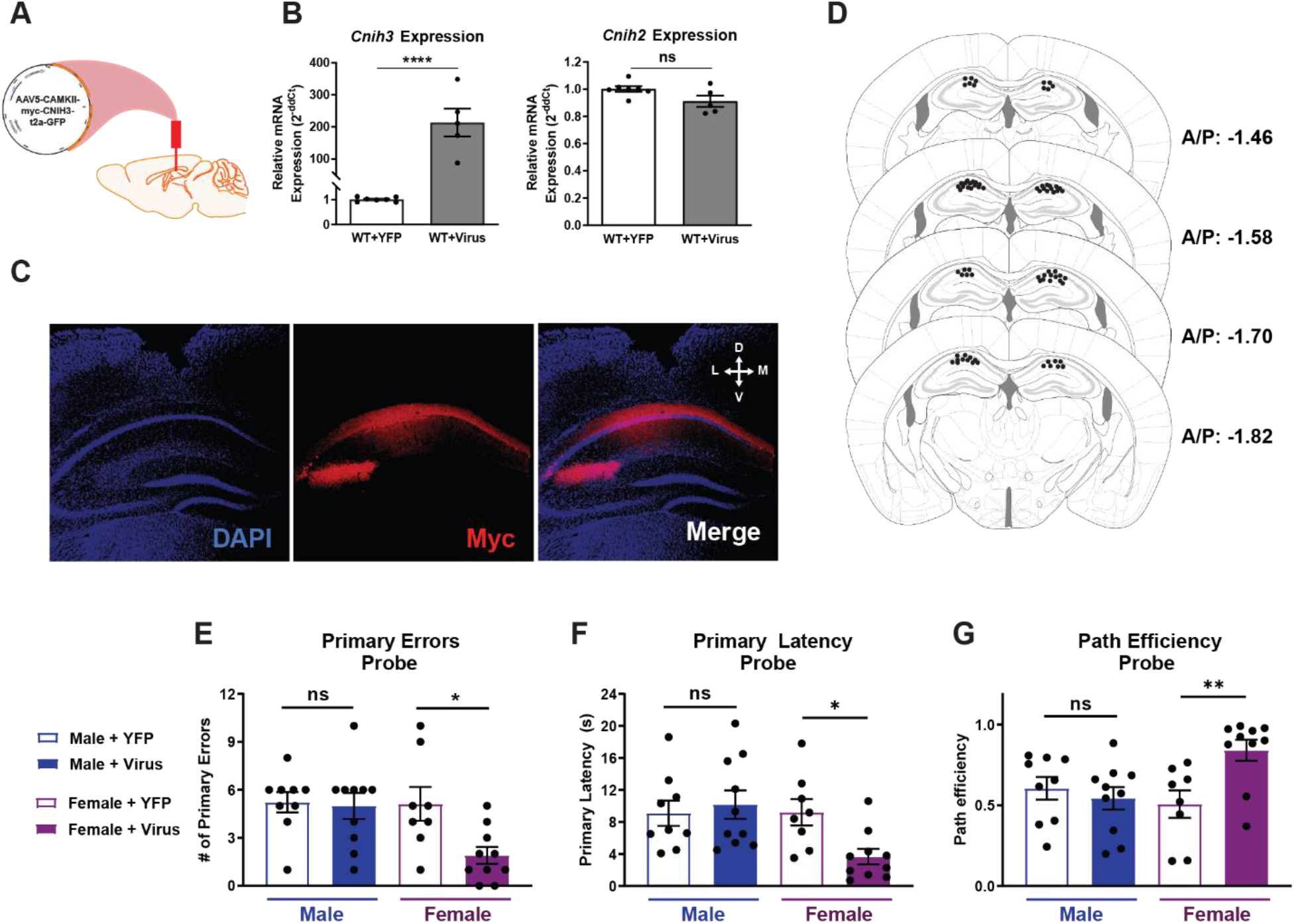
Spatial memory is enhanced in female mice overexpressing *Cnih3* in the dorsal hippocampus. (A) AAV5-CAMKII-myc-CNIH3-t2a-GFP virus was injected bilaterally into the dorsal hippocampus of WT male and female mice to induce the overexpression of *Cnih3* (−1.8 mm A/P, ±1.4 mm M/L, −1.8 mm D/V). (B) RT-qPCR analysis of *Cnih3* and *Cnih2* mRNA in the dorsal hippocampus 3 weeks post viral injection of YFP control (n = 7) and *Cnih3* overexpression (n = 5) viruses (two-tailed unpaired *t*-test; ns *p* > 0.05, **** *p* < 0.0001). (C) Representative image of the dorsal hippocampus from a brain injected with AAV5-CAMKII-myc-CNIH3-t2a-GFP. Brain slices were probed for the myc-tag adjacent to *Cnih3* in the viral construct. (D) Injection placements for AAV5-CAMKII-myc-CNIH3-t2a-GFP and control AAV5-CAMKII-eYFP viruses of animals tested in the Barnes maze. (E – G) Barnes maze results of male and female mice injected with either control (male: n = 9; female: n = 8) or *Cnih3* overexpression (male: n = 10; female: n = 10) virus during the probe trails is represented by (E) primary errors before location of the target hole, (F) primary latency to entry into the target hole, and (G) path efficiency to the target hole (two-way ANOVA for genotype and sex and *post-hoc* Sidak’s multiple comparisons test; ns *p* > 0.05, * *p* < 0.05, ** *p* < 0.01).

### Spatial memory is improved in female mice overexpressing Cnih3 in the dorsal hippocampus

To determine if *Cnih3* overexpression localized in the dorsal hippocampus was sufficient to enhance spatial memory, *Cnih3* was overexpressed in the dorsal hippocampus of WT male and female mice prior to spatial memory testing in the Barnes Maze (Figure 4C-D). During the Day 5 probe, we observed significant main effects of sex and viral expression on primary errors (two-way ANOVA, sex [F_(1, 33)_ = 4.362, *p* = 0.0445], virus [F_(1, 33)_ = 5.071, *p* = 0.0311], and interaction [F_(1, 33)_ = 3.848, *p* = 0.0583]) (Figure 4E), and significant main effects of sex for primary latency (sex [F_(1, 33)_ = 4.429, *p* = 0.0430], virus [F_(1, 33)_ = 4.429, *p* = 0.1497], and interaction [F_(1, 33)_ = 4.783, *p* = 0.0359]) (Figure 4F). Sex and viral expression did not significantly affect path efficiency in the Barnes maze, but a significant interaction between these two effects was observed (sex [F_(1, 33)_ = 1.908, *p* = 0.1765], virus [F_(1, 33)_ = 3.536, *p* = 0.0689], and interaction [F_(1, 33)_ = 7.438, *p* = 0.0101]) (Figure 4G). To investigate the sex-dependent effect of viral expression on path efficiency within our non-normal dataset, a nonparametric Mann-Whitney two-tailed test was also utilized to identify a significant effect of *Cnih3* hippocampal overexpression on path efficiency in the Barnes maze in female mice (Female, *p* = 0.0062; Male, *p* = 0.4002). Overall, we conclude that *Cnih3* overexpression in the dorsal hippocampus significantly affects performance in the Barnes maze in a sex-specific manner.

To further dissect the interaction between sex and *Cnih3* hippocampal overexpression, *post-hoc* multiple comparison analysis was conducted to compare each dataset to YFP expressing control mice. Female mice overexpressing *Cnih3* in the hippocampus committed fewer primary errors (*post-hoc* Sidak’s multiple comparisons test, *p* = 0.0121) (Figure 4E), took less time to reach the target (*p* = 0.0310) (Figure 4F), and took a more efficient path to the target (*p* = 0.0059) (Figure 4G) compared to control females. Male mice overexpressing *Cnih3* in the hippocampus demonstrated no significant difference in the number of primary errors (*p* = 0.9731) (Figure 4F), primary latency (*p* = 0.8497) (Figure 4G), nor path efficiency (*p* = 0.7947) (Figure 4G) compared to male mice expressing the control virus in the hippocampus. We did not observe any significant differences between YFP expressing control male and female mice in the number of primary errors (*p* = 0.9953), the primary latency (*p* = 0.9980), nor the path efficiency (*p* = 0.5993). Overall, female mice overexpressing *Cnih3* in the dorsal hippocampus exhibited enhanced spatial memory in the Barnes maze compared to YFP expressing female controls.

## DISCUSSION

We have developed and characterized a line of *Cnih3*^+/+^, *Cnih3*^+/-^, and *Cnih3*^-/-^ C57BL/6 mice to investigate the role of CNIH3 in the formation of spatial memory. We observed a significant attenuation of spatial memory in *Cnih3*^-/-^ female mice compared to sex-matched controls, and a congruous enhancement of spatial memory only in female mice overexpressing *Cnih3* in the dorsal hippocampus. Therefore, we have identified novel sex differences in the function of CNIH3 in the brain.

While prior studies have not investigated the role of sex in the function of AMPAR auxiliary proteins, studies of sex differences in AMPAR-dependent spatial memory tasks offer mixed results. Some studies reported overall superior spatial memory in WT male compared to female rodents (Monfort et al., 2015); others observed sex differences in search strategies, but not performance in spatial memory tasks (Locklear and Kritzer, 2014). Additional studies have reported sex differences only in spatial memory retention but not in short-term memory (Qi et al., 2016) or have found no spatial memory differences due to sex in WT young adult rodents (Frick et al., 1999; Dachtler et al., 2011). We did not observe differences in spatial memory in the Barnes maze between WT male and female 8 – 12-week-old mice. However, due to the clear effect of sex on CNIH3-dependent spatial memory, future studies of CNIH3 will monitor female estrous cycles to control for the potential effects of estrous on AMPARs and spatial memory.

Multiple studies have also shown a sex-specific component in AMPAR-mediated synaptic plasticity underlying memory. Several studies have reported that differences between male and female rats in evoked AMPAR/NMDAR signaling in hippocampal synapses, along with differences in the magnitude of evoked LTP, may underlie sex differences in spatial memory (Monfort et al., 2015; Qi et al., 2016). Estrous has been shown to modulate diffusion of AMPARs to the surface in female mice (Palomero-Gallagher et al., 2003; Tada et al., 2015; Bechard et al., 2018), which may underlie part of the reported importance of estrogenic mechanisms for memory in females (Cordeira et al., 2018; Frick et al., 2018; Koss et al., 2018; Wang et al., 2018; Koebele et al., 2019). However, the sex-specific role of AMPARs in memory may also be specific to the type of memory task being tested, as a study by Dachtler *et al*. concluded that the GluA1 subunit was necessary only for male, not female, mice in fear conditioning memory, but that hippocampal GluA1 was necessary for spatial learning in both sexes (Dachtler et al., 2011). Therefore, more investigation is needed to determine how AMPARs and their auxiliary proteins mediate sexspecific spatial memory mechanisms.

Furthermore, we are particularly interested in extending our findings to examine the role of CNIH3 in contextual opioid-associated memory. We previously reported an association between SNPs in CNIH3 and protection from opioid dependence in humans (Nelson et al., 2016). In addition, we have previously shown the importance of the dorsal hippocampus (Fakira et al., 2016; Williams et al., 2019) and AMPAR signaling in the modulation of opioid-associated contextual memory (Morón et al., 2007; Billa et al., 2009; Billa et al., 2010; Xia et al., 2011). Sex differences in AMPAR activity have also been linked to cocaine-associated memory (Bechard et al., 2018; Ganguly et al., 2019). Therefore, the novel component of sex differences in spatial memory identified here suggest that CNIH3 could play a role in sex-dependent drug-associated memory.

In conclusion, we have developed a line of *Cnih3* KO mice and characterized the effect of *Cnih3* expression on spatial memory in mice. Despite no differences in spatial memory observed between WT male and female mice, only female *Cnih3^-/-^* mice exhibited attenuated spatial memory, whereas female *Cnih3* overexpressing mice exhibited an enhancement of spatial memory. This was the first study to identify a sex difference in the function of CNIH3, or in any AMPAR auxiliary protein to our knowledge. The results of this study offer insight into sex-dependent AMPAR auxiliary protein regulation of memory, which may impact a wide range of AMPAR-dependent behaviors and disorders.

## Acknowledgements

This work was funded by grants from the National Institutes of Health (NIH) R21/R33-DA041883 to ECN, JDD, and JAM, R01-DA042499 to JAM, and a Washington University Program in Neuroscience Training Grant (T32-GM008151) to HEF. The authors would also like to thank Dr. Rebecca Ouwenga for her technical assistance.

## REFERENCES

Barnes CA (1979) Memory deficits associated with senescence: a neurophysiological and behavioral study in the rat. J Comp Physiol Psychol 93:74–104.

Bechard AR, Hamor PU, Schwendt M, Knackstedt LA (2018) The effects of ceftriaxone on cue-primed reinstatement of cocaine-seeking in male and female rats: estrous cycle effects on behavior and protein expression in the nucleus accumbens. Psychopharmacology (Berl) 235:837–848.

Billa SK, Sinha N, Rudrabhatla SR, Morón JA (2009) Extinction of morphine-dependent conditioned behavior is associated with increased phosphorylation of the GluR1 subunit of AMPA receptors at hippocampal synapses. Eur J Neurosci 29:55–64.

Billa SK, Liu J, Bjorklund NL, Sinha N, Fu Y, Shinnick-Gallagher P, Morón JA (2010) Increased insertion of glutamate receptor 2-lacking alpha-amino-3-hydroxy-5-methyl-4-isoxazole propionic acid (AMPA) receptors at hippocampal synapses upon repeated morphine administration. Mol Pharmacol 77:874–883.

Boudkkazi S, Brechet A, Schwenk J, Fakler B (2014) Cornichon2 dictates the time course of excitatory transmission at individual hippocampal synapses. Neuron 82:848–858.

Brockie PJ, Jensen M, Mellem JE, Jensen E, Yamasaki T, Wang R, Maxfield D, Thacker C, Hoerndli F, Dunn PJ, Tomita S, Madsen DM, Maricq AV (2013) Cornichons control ER export of AMPA receptors to regulate synaptic excitability. Neuron 80:129–142.

Brown P, McGuire H, Bowie D (2018) Stargazin and cornichon-3 relieve polyamine block of AMPA receptors by enhancing blocker permeation. J Gen Physiol 150:67–82.

Challenor M, O’Hare Doig R, Fuller P, Giacci M, Bartlett C, Wale CH, Cozens GS, Hool L, Dunlop S, Swaminathan Iyer K, Rodger J, Fitzgerald M (2015) Prolonged glutamate excitotoxicity increases GluR1 immunoreactivity but decreases mRNA of GluR1 and associated regulatory proteins in dissociated rat retinae in vitro. Biochimie 112:160–171.

Cheng J, Dong J, Cui Y, Wang L, Wu B, Zhang C (2012) Interacting partners of AMPA-type glutamate receptors. J Mol Neurosci 48:441–447.

Coombs ID, Soto D, Zonouzi M, Renzi M, Shelley C, Farrant M, Cull-Candy SG (2012) Cornichons modify channel properties of recombinant and glial AMPA receptors. J Neurosci 32:9796–9804.

Cordeira J, Kolluru SS, Rosenblatt H, Kry J, Strecker RE, McCarley RW (2018) Learning and memory are impaired in the object recognition task during metestrus/diestrus and after sleep deprivation. Behavioural Brain Research 339:124–129.

Dachtler J, Fox KD, Good MA (2011) Gender specific requirement of GluR1 receptors in contextual conditioning but not spatial learning. Neurobiol Learn Mem 96:461–467.

Drummond JB, Simmons M, Haroutunian V, Meador-Woodruff JH (2012) Upregulation of cornichon transcripts in the dorsolateral prefrontal cortex in schizophrenia. Neuroreport 23:1031–1034.

Fakira AK, Massaly N, Cohensedgh O, Berman A, Morón JA (2016) Morphine-Associated Contextual Cues Induce Structural Plasticity in Hippocampal CA1 Pyramidal Neurons. Neuropsychopharmacology 41:2668–2678.

Floor K, Barøy T, Misceo D, Kanavin OJ, Fannemel M, Frengen E (2012) A 1 Mb de novo deletion within 11q13.1q13.2 in a boy with mild intellectual disability and minor dysmorphic features. Eur J Med Genet 55:695–699.

Frick KM, Burlingame LA, Arters JA, Berger-Sweeney J (1999) Reference memory, anxiety and estrous cyclicity in C57BL/6NIA mice are affected by age and sex. Neuroscience 95:293–307.

Frick KM, Tuscher JJ, Koss WA, Kim J, Taxier LR (2018) Estrogenic regulation of memory consolidation: A look beyond the hippocampus, ovaries, and females. Physiol Behav 187:57–66.

Gandhi RM, Kogan CS, Messier C, Macleod LS (2014) Visual-spatial learning impairments are associated with hippocampal PSD-95 protein dysregulation in a mouse model of fragile X syndrome. Neuroreport 25:255–261.

Ganguly P, Honeycutt JA, Rowe JR, Demaestri C, Brenhouse HC (2019) Effects of early life stress on cocaine conditioning and AMPA receptor composition are sex-specific and driven by TNF. Brain Behav Immun 78:41–51.

Haering SC, Tapken D, Pahl S, Hollmann M (2014) Auxiliary subunits: shepherding AMPA receptors to the plasma membrane. Membranes (Basel) 4:469–490.

Harmel N, Cokic B, Zolles G, Berkefeld H, Mauric V, Fakler B, Stein V, Klöcker N (2012) AMPA receptors commandeer an ancient cargo exporter for use as an auxiliary subunit for signaling. PLoS One 7:e30681.

Herring BE, Shi Y, Suh YH, Zheng CY, Blankenship SM, Roche KW, Nicoll RA (2013) Cornichon proteins determine the subunit composition of synaptic AMPA receptors. Neuron 77:1083–1096.

IMPC (2016) Cnih3tm1a(KOMP)Wtsi. In: (Consortium IMP, ed).

Kato AS, Gill MB, Ho MT, Yu H, Tu Y, Siuda ER, Wang H, Qian YW, Nisenbaum ES, Tomita S, Bredt DS (2010) Hippocampal AMPA receptor gating controlled by both TARP and cornichon proteins. Neuron 68:1082–1096.

Kent WJ, Sugnet CW, Furey TS, Roskin KM, Pringle TH, Zahler AM, Haussler aD (2002) The Human Genome Browser at UCSC. Genome Research 12:996–1006.

Kim S, Thompson RF (2011) c-Fos, Arc, and stargazin expression in rat eyeblink conditioning. Behav Neurosci 125:117–123.

Koebele SV, Palmer JM, Hadder B, Melikian R, Fox C, Strouse IM, DeNardo DF, George C, Daunis E, Nimer A, Mayer LP, Dyer CA, Bimonte-Nelson HA (2019) Hysterectomy Uniquely Impacts Spatial Memory in a Rat Model: A Role for the Nonpregnant Uterus in Cognitive Processes. Endocrinology 160:1–19.

Koss WA, Haertel JM, Philippi SM, Frick KM (2018) Sex Differences in the Rapid Cell Signaling Mechanisms Underlying the Memory-Enhancing Effects of 17β-Estradiol. eNeuro 5.

Lee HK, Takamiya K, Han JS, Man H, Kim CH, Rumbaugh G, Yu S, Ding L, He C, Petralia RS, Wenthold RJ, Gallagher M, Huganir RL (2003) Phosphorylation of the AMPA receptor GluR1 subunit is required for synaptic plasticity and retention of spatial memory. Cell 112:631–643.

Li J, Han Z, Cao B, Cai CY, Lin YH, Li F, Wu HY, Chang L, Luo CX, Zhu DY (2017) Disrupting nNOS-PSD-95 coupling in the hippocampal dentate gyrus promotes extinction memory retrieval. Biochem Biophys Res Commun 493:862–868.

Locklear MN, Kritzer MF (2014) Assessment of the effects of sex and sex hormones on spatial cognition in adult rats using the Barnes maze. Horm Behav 66:298–308.

Matsuo N, Reijmers L, Mayford M (2008) Spine-type-specific recruitment of newly synthesized AMPA receptors with learning. Science 319:1104–1107.

Mauric V, Möders A, Harmel N, Heimrich B, Sergeeva OA, Klöcker N (2013) Ontogeny repeats the phylogenetic recruitment of the cargo exporter cornichon into AMPA receptor signaling complexes. Mol Cell Neurosci 56:10–17.

Monfort P, Gomez-Gimenez B, Llansola M, Felipo V (2015) Gender differences in spatial learning, synaptic activity, and long-term potentiation in the hippocampus in rats: molecular mechanisms. ACS Chem Neurosci 6:1420–1427.

Morón JA, Abul-Husn NS, Rozenfeld R, Dolios G, Wang R, Devi LA (2007) Morphine administration alters the profile of hippocampal postsynaptic density-associated proteins: a proteomics study focusing on endocytic proteins. Mol Cell Proteomics 6:29–42.

Nagura H, Ishikawa Y, Kobayashi K, Takao K, Tanaka T, Nishikawa K, Tamura H, Shiosaka S, Suzuki H, Miyakawa T, Fujiyoshi Y, Doi T (2012) Impaired synaptic clustering of postsynaptic density proteins and altered signal transmission in hippocampal neurons, and disrupted learning behavior in PDZ1 and PDZ2 ligand binding-deficient PSD-95 knockin mice. Molecular Brain 5:43.

Nelson EC et al. (2016) Evidence of CNIH3 involvement in opioid dependence. Mol Psychiatry 21:608–614.

Palomero-Gallagher N, Bidmon HJ, Zilles K (2003) AMPA, kainate, and NMDA receptor densities in the hippocampus of untreated male rats and females in estrus and diestrus. J Comp Neurol 459:468–474.

Park J, Chávez AE, Mineur YS, Morimoto-Tomita M, Lutzu S, Kim KS, Picciotto MR, Castillo PE, Tomita S (2016) CaMKII Phosphorylation of TARPγ-8 Is a Mediator of LTP and Learning and Memory. Neuron 92:75–83.

Penn AC, Zhang CL, Georges F, Royer L, Breillat C, Hosy E, Petersen JD, Humeau Y, Choquet D (2017) Hippocampal LTP and contextual learning require surface diffusion of AMPA receptors. Nature 549:384–388.

Qi X, Zhang K, Xu T, Yamaki VN, Wei Z, Huang M, Rose GM, Cai X (2016) Sex Differences in Long-Term Potentiation at Temporoammonic-CA1 Synapses: Potential Implications for Memory Consolidation. PLoS One 11:e0165891.

Sanderson DJ, Good MA, Seeburg PH, Sprengel R, Rawlins JN, Bannerman DM (2008) The role of the GluR-A (GluR1) AMPA receptor subunit in learning and memory. Prog Brain Res 169:159–178.

Sebastian V, Estil JB, Chen D, Schrott LM, Serrano PA (2013) Acute physiological stress promotes clustering of synaptic markers and alters spine morphology in the hippocampus. PLoS One 8:e79077.

Shanks NF, Cais O, Maruo T, Savas JN, Zaika EI, Azumaya CM, Yates JR, Greger I, Nakagawa T (2014) Molecular dissection of the interaction between the AMPA receptor and cornichon homolog-3. J Neurosci 34:12104–12120.

Shi Y, Suh YH, Milstein AD, Isozaki K, Schmid SM, Roche KW, Nicoll RA (2010) Functional comparison of the effects of TARPs and cornichons on AMPA receptor trafficking and gating. Proc Natl Acad Sci U S A 107:16315–16319.

Sunyer B, Patil S, Höger H, Lubec G (2007) Barnes Maze, a useful task to assess spatial reference memory in the mice. Protocol Exchange.

Tada H, Koide M, Ara W, Shibata Y, Funabashi T, Suyama K, Goto T, Takahashi T (2015) Estrous Cycle-Dependent Phasic Changes in the Stoichiometry of Hippocampal Synaptic AMPA Receptors in Rats. PLoS One 10:e0131359.

Torquatto KI, Menegolla AP, Popik B, Casagrande MA, de Oliveira Alvares L (2019) Role of calcium-permeable AMPA receptors in memory consolidation, retrieval and updating. Neuropharmacology 144:312–318.

Trifonov S, Yamashita Y, Kase M, Maruyama M, Sugimoto T (2016) Overview and assessment of the histochemical methods and reagents for the detection of β-galactosidase activity in transgenic animals. Anat Sci Int 91:56–67.

Tzakis N, Bosnic T, Ritchie T, Dixon K, Holahan MR (2016) The effect of AMPA receptor blockade on spatial information acquisition, consolidation and expression in juvenile rats. Neurobiol Learn Mem 133:145–156.

Volk L, Kim C-H, Takamiya K, Yu Y, Huganir RL (2010) Developmental regulation of protein interacting with C kinase 1 (PICK1) function in hippocampal synaptic plasticity and learning.

Wang W, Le AA, Hou B, Lauterborn JC, Cox CD, Levin ER, Lynch G, Gall CM (2018) Memory-Related Synaptic Plasticity Is Sexually Dimorphic in Rodent Hippocampus. J Neurosci 38:7935–7951.

West DB, Pasumarthi RK, Baridon B, Djan E, Trainor A, Griffey SM, Engelhard EK, Rapp J, Li B, de Jong PJ, Lloyd KC (2015) A lacZ reporter gene expression atlas for 313 adult KOMP mutant mouse lines. Genome Res 25:598–607.

Williams SB, Arriaga M, Post WW, Korgaonkar AA, Moron JA, Han EB (2019) Hippocampal Activity Dynamics During Contextual Reward Association in Virtual Reality Place Conditioning. bioRxiv:545608.

Xia Y, Portugal GS, Fakira AK, Melyan Z, Neve R, Lee HT, Russo SJ, Liu J, Morón JA (2011) Hippocampal GluA1-containing AMPA receptors mediate context-dependent sensitization to morphine. J Neurosci 31:16279–16291.

